# Stretching DNA to twice the normal length with single-molecule hydrodynamic trapping

**DOI:** 10.1101/801464

**Authors:** Yan Jiang, Theodore Feldman, Julia A.M. Bakx, Darren Yang, Wesley P. Wong

## Abstract

Single-molecule force spectroscopy has brought many new insights into nanoscale biology, from the functioning of molecular motors, to the mechanical response of soft materials within the cell. To expand the single-molecule toolbox, we have developed a surface-free force spectroscopy assay based on a high-speed hydrodynamic trap capable of applying extremely high tensions for long periods of time. High-speed single-molecule trapping is enabled by a rigid and gas-impermeable microfluidic chip, rapidly and inexpensively fabricated out of glass, double-sided tape and UV-curable adhesive. Our approach does not require difficult covalent attachment chemistries, and enables simultaneous force application and single-molecule fluorescence. Using this approach, we have induced a highly extended state with twice the contour length of B-DNA in regions of partially intercalated double-stranded (dsDNA) by applying forces up to 250 pN. This highly extended state resembles the hyperstretched state of dsDNA, which was initially discovered as a structure fully intercalated by dyes under high tension. It has been hypothesized that hyperstretched DNA could also be induced without the aid of intercalators if high-enough forces were applied, which matches our observation. Combining force application with single-molecule fluorescence imaging is critical for distinguishing hyperstretched DNA from single-stranded DNA that can result from peeling. High-speed hydrodynamic trapping is a powerful yet accessible force spectroscopy method that enables the mechanics of biomolecules to be probed in previously difficult to access regimes.

## Introduction

Single-molecule force spectroscopy has greatly advanced our understanding of macromolecules, illuminating the mechanical properties of nucleic acids, the dynamics of motor proteins, and the kinetics of receptor-ligand binding^1^. Although current approaches for force spectroscopy are powerful and relatively mature, the surface attachments required by most force spectroscopy methods^1–8^ often impose challenges. In particular, high force single-molecule measurements, for example in the range above 150 pN, can be difficult to perform in the equilibrium regime, because many common surface attachments strategies (e.g biotin-streptavidin or digoxigenin-antidigoxigenin) have limited lifetimes and are rapidly ruptured at constant high forces^9–13^. Even with multiple biotin-streptavidin bonds, the maximum force reported for near equilibrium conditions on a time scale of 1-10 minutes is ∼ 180 pN^14,15^. In some cases, parts of the molecule can strongly adsorb to gold surfaces and AFM tips, allowing constant application of hundreds of pN^16^. When physical adsorption is unavailable or undesirable, more involved covalent conjugation chemistries are needed for high force application^13,17–23^. Furthermore, the high force regime is outside the practical range of many standard approaches—for example, optical tweezers can run into problems with available laser power, local heating, or free radical generation, and magnetic tweezers typically do not have strong enough field gradients to generate these forces with standard magnetic microspheres^1,24,25^. These problems limit investigations in the high force regime under conditions of constant force or loading rates slow enough to explore near-equilibrium behaviour. In addition, specific and homogeneous surface attachment can often be challenging to carry out, and can potentially lead to artifacts due to denaturation or distortion of molecules at the surface, or the addition of colloidal forces (e.g. Van der Waals) that may be undesirable^26–28^.

Here, we present a surface-free force spectroscopy approach that is not only inexpensive and accessible, but also able to probe tensions in the very high force regime without the need for challenging attachment chemistries. This method is based on an active-feedback hydrodynamic trap for particles and cells, pioneered by Schroeder *et al*^29–31^ using a cross-slot flow channel^32^. Flow gradients, such as shear and elongational flow ^31–36^, have been used to induce tension in untethered polymers, but single-molecule observation and force application is usually limited to a low level or a short amount of time as the molecule can simply flow away from the region of interest. Tanyeri *et al*^29,37,38^ tackled this problem by combining elongational flow with automated active feedback of the outlet pressure using dual layer PDMS microfluidics. The hydrodynamic trap has been further developed to enable two-dimensional trapping, tighter trapping, and trapping of multiple particles by implementing multiple feedback channels, single-layer device design, and model predictive control (MPC) algorithms^37,39–42^ The hydrodynamic trap and the newer Stokes trap have been used to measure the properties of complex fluids and to study the dynamics of single polymers, including DNA, under elongational flow^43–47^. We have developed a simple and rapid approach to fabricate microfluidic devices for fluorescence-based hydrodynamic trapping, optimal for single-molecule force spectroscopy applications. Using this device, we have trapped single DNA molecules under high elongation rates and have induced extremely high tensions in them. The flow elongation rate is defined as the gradient of flow velocity along the direction of flow. Our microfluidic chip is rapidly and inexpensively fabricated out of glass, die cut double-sided tape and UV-cured adhesive. It is rigid and gas impermeable, ideal for high-pressure and fluorescence imaging applications. Our high-speed hydrodynamic trap can trap and apply high forces to single polymers or filaments under elongational flow, enabling us to perform force spectroscopy over a wide range of forces in solution without tethering. We have demonstrated the application of controlled tensions up to 250pN on a single T4 DNA molecule without requiring the attachment of any beads or surfaces. High-speed hydrodynamic trapping can serve as a powerful yet accessible force spectroscopy method, particularly in the high force regime. Furthermore, our approach enables simultaneous monitoring of the sample with single-molecule fluorescence, which can provide essential information about the conformations and functions of biomolecules.

Mechanical studies of DNA have served as a corner stone for single-molecule measurements, and have led to new insights into a range of biological processes^48–57^. DNA has served not only as a subject of study, but also as a tool for the creation of handles and linkages with mechanical signatures, enabling the characterization of a variety of molecular structures and functions^58–62^. More generally, the field of structural DNA nanotechnology has established DNA as a versatile programmable material for the creation of complex nanoscale structures and molecular devices ^63,64^. Therefore, a more comprehensive understanding of DNA mechanics would be of great value to both mechanobiology, and to DNA nanotechnology.

dsDNA exhibits complex elastic properties due to its characteristic base stacking and double-helical structure^65^. For example, B-DNA, the structure most commonly found in nature, can transition to different structures under mechanical force. It has been known for over two decades that above a critical overstretching force around 65pN, B-DNA undergoes a transition to reach a contour length of 0.58 nm/bp, 70% longer than the 0.34 nm/bp of B-DNA ^5,66^, and it has been established that this can occur in the absence of peeling^67–69^. Recently, a new state of dsDNA with an even longer contour length of 0.7 nm/bp—close to the maximum length allowed by the standard bond lengths within the backbone of DNA—was discovered under high force in the presence of intercalating dyes^14^. Accordingly, they called this conformation hyperstretched DNA. Using fluorescently stained single DNA molecules, they were able to confirm that the DNA remained hybridized, and peeling did not occur, and also noted that it was fully intercalated by dyes, with the next-neighbour exclusion rule^70^ overcome by force. The authors proposed that hyperstretched DNA may be induced on naked DNA without intercalators in the extremely high force regime, yet testing this hypothesis is extremely challenging with standard experimental approaches. Using our approach, we have stretched regions of partially dye intercalated double-stranded DNA (dsDNA) to twice its normal length by applying forces up to 250 pN. The preserved intercalating dye fluorescence suggests that the two DNA strands remained hybridized. Our result supports the hypothesis that force alone is capable of inducing this hyperstretched state of DNA.

## Experimental

### Microfluidic cross-slot chip

To construct the microfluidic flow channel, detergent cleaned #1.5 cover glass (Gold Seal, Thermo Scientific, Waltham, MA, USA) was overlaid with a central piece of 40 μm thick, double-sided polyester tape (ARcare 92712, Adhesive Research, Glen Rock, PA, USA) with a cut-out pattern shown in Supplementary Figure 1 and topped with a 1.1 mm thick glass slide (VWR, USA) with two access holes for each channel. To clean the #1.5 cover glasses, they were immersed in 1% (v/v) Hellmanex III solution (Hellma, Müllheim, Germany), heated to about 80 °C, sonicated for 1 min, and rinsed with Millipore water before being assembled for flow channels. The pattern was cut out from the tape using a die cutter (CE3000, Graphtec America, Irvine, CA, USA) fitted with a 60° angle blade (AGTK-60, Clean Cut Blade, Waterloo, IA, USA), following the sequence shown in Supplementary Figure 1. The pattern can also be cut with an Excimer laser, which produces more accurate lines. But the equipment is more expensive and the procedure is time-consuming. The performance of the die cutter is sufficient for this application. After the glass-tape-glass sandwich was assembled, the adhesive channels were filled with optical adhesive (NOA81, Norland Products, Cranbury, NJ, USA) using vacuum suction. The optical adhesive was cured by 365 nm UV illumination. The chip was then baked in a 100 °C oven for 1 hour or autoclaved to strengthen the bonding and smoothen the edges of the channels. A ∼3 mm thick polydimethylsiloxane slab with 0.75 mm diameter holes was aligned with the access holes and clamped onto the glass sandwich. To avoid trapping air bubbles in the PDMS holes, the PDMS surface was made hydrophilic via oxygen plasma activation prior to assembly. The holes on each end of the flow channel were connected through 1/32” outer diameter PEEK tubing to buffer or sample. The buffer inlets and the feedback outlet were connected via 0.01” and 0.02” inner diameter PEEK tubing (1569, IDEX Health & Science, Middleborough, MA, USA) to buffer in 4 ml gas tight vials containing nitrogen gas. The pressures of the nitrogen gas in the two bottles were controlled by a high-precision electronic pressure regulator (DQPV1TFEE010CXL, Proportion-Air, McCordsville, IN, USA) connected to a compressed nitrogen cylinder (>99% pure, Lifegas, Marlborough, MA, USA) to control the flow rate and to provide feedback, respectively. The other outlet was connected to buffer in an open container placed ∼ 20 cm above the chip to provide a constant pressure. The sample inlets were connected to two 25 μl glass syringes via 0.02” inner diameter PEEK tubing.

### Fluorescence imaging and active feedback

The fluorescence microscopy system was custom built on a vibration damping optical table (Thorlabs) with a 60× oil immersion objective (NA 1.49, CFI Apo TIRF 60× H, Nikon, Japan), 485 nm laser (CUBE 485-30C, Coherent, Santa Clara, CA, USA), with an oscillating diffuser (laser speckle reducer; Optotune, Switzerland), and an EMCCD camera (C9100, Hamamatsu, Japan). Synchronized image recording and flow was controlled with custom software (LabView, National Instruments, Austin, TX, USA).

When a fluorescent molecule flowed into the cross slot, each image frame was analysed in real-time by the custom LabView code to determine the centre of the molecule. A feedback voltage was calculated based on the vertical position of the molecule using a PID controller and applied to the electronic pressure regulator. The PID parameters were empirically selected to optimize the trapping performance. The fluorescent beads used for characterizing trap performance were FluoSpheres microspheres (F8827, ThermoFisher, Waltham, MA, USA). They were trapped in a 12.6 centipoise (cP) pH7.4 1xPBS buffer containing 137 mM NaCl, 10 mM Phosphate, 2.7 mM KCl and 48 % weight/weight sucrose.

### Labelling of DNA

2 μl of 5 ng/μl T4 GT7 DNA (318-03971, Nippon Gene, Tokyo, Japan) was slowly added to 298 μl diluted YOYO-1 solution to reach a final concentration of 52 nM DNA base pairs and 26 or 10.4 nM YOYO-1, for a 2:1 or 5:1 base pair to dye ratio. Wide bore pipet tips were used to handle T4 DNA to avoid damaging them by shear flow. DNA and dye solution were mixed by slowly stirring with a pipet tip. The 298 μl YOYO-1 solution was prepared by diluting 0.98 or 0.39 μL 1 mM YOYO-1 (Y3601, ThermoFisher, Waltham, MA, USA) into 297.0 or 297.6 μl TE buffer. The intercalating of each YOYO-1 molecule increases the contour length of DNA by 0.34 nm but does not change the persistent length of DNA^71,72^. For 2:1 stained T4 DNA, the contour length was ∼ 0.43 nm/bp when in the partially intercalated B-DNA form. This means 26% of the spaces between base pairs were intercalated since intercalated base pairs result in a contour length of 0.68 nm/bp, corresponding to a nucleic acid base pairs to dye monomer ratio (N:D) of about 4:1. For 5:1 stained DNA, the contour length was ∼ 0.38 nm/bp when in the partially intercalated B-DNA form. Given that each intercalated monomer increases the contour length by 0.34 nm, ∼12% of the spaces between base pairs were intercalated, corresponding to a nucleic acid base pairs to dye monomer ratio (N:D) of about 8:1. The DNA/dye mixtures were then incubated at 50°C for 2 hours^73^ and stored at 4°C until use.

### Trapping and stretching of DNA

Buffer used for the trapping experiments (trapping buffer) contained 60% weight/weight sucrose, 10 mM Tris-HCL pH7.5, 140 mM 2-mercaptoethanol (BME), 3.5 mM protocatechuic acid (PCA, Santa Cruz Biotechnology, Dallas, Texas, USA), 20 nM protocatechuate-3,4-dioxygenase (PCD, P87279, Sigma-Aldrich, St. Louis, Missouri, USA). The combination of the gas-impermeable microfluidic chip and the oxygen scavenger system greatly reduced photobleaching of the dyes and photo-induced damage to DNA. To reduce shear damage to the DNA, the DNA sample was slowly loaded (0.5 μl/min) into the PEEK tubing connected to the left syringe before the PEEK tubing was connected to the chip. To start the experiments, DNA was injected into the cross-slot from the left sample injection port at 0.5-1 μl/hr (main text Figure 1D) and the buffer was injected into the buffer inlets at flow rates that induced 0-5/s elongational flow in the cross-slot. Upon the capture of a DNA molecule, the sample injection was turned off to prevent more molecules from entering the cross-slot. Meanwhile, the buffer injection rate was adjusted to achieve the specified flow elongation rate.

**Figure 1.**
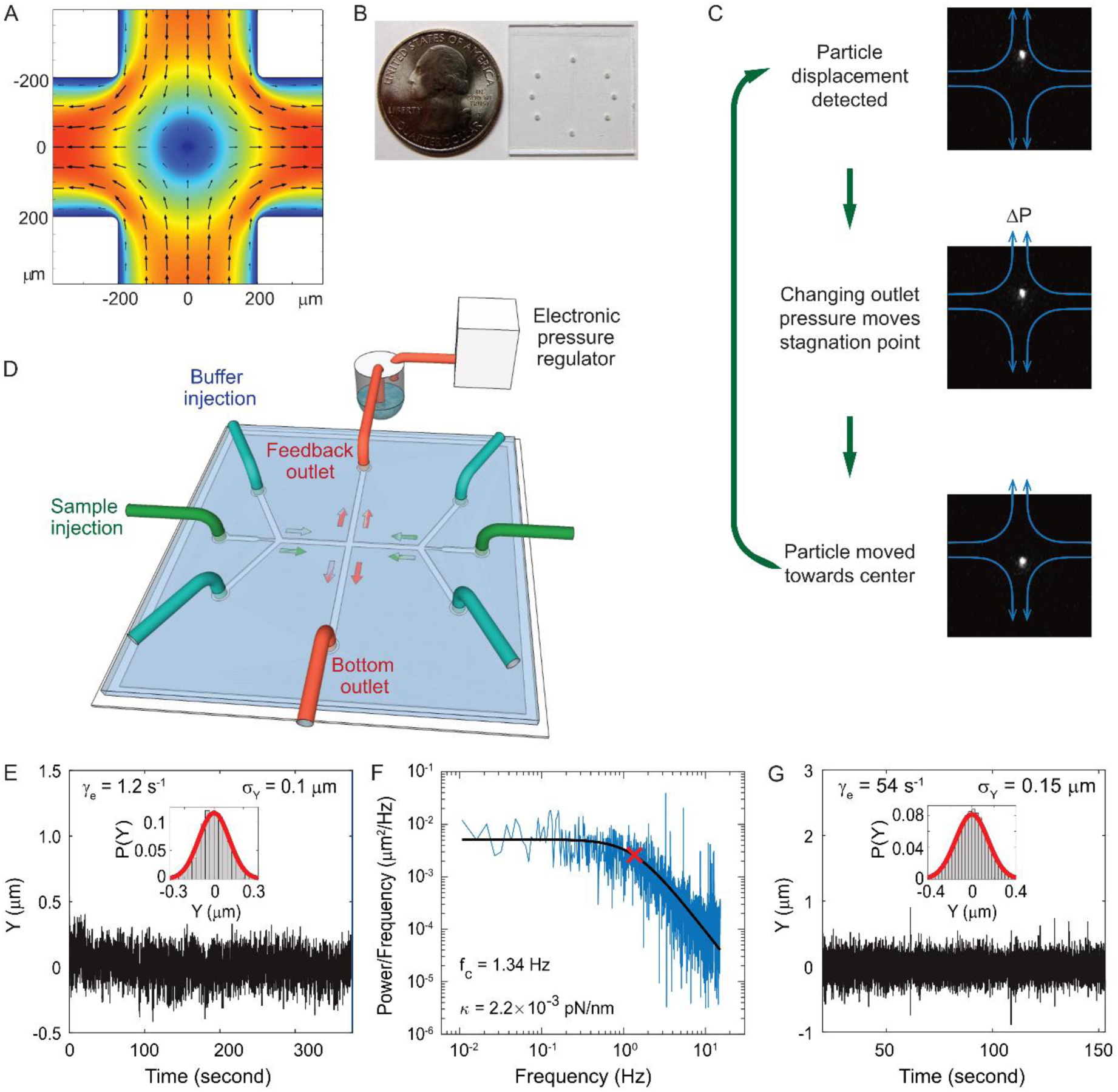
High-speed active-feedback hydrodynamic trap design and performance. A) Elongational flow in cross slot. Small arrows indicate the flow direction. Colour indicates the flow velocity with red being the fastest and blue the slowest. B) A photo of an actual hydrodynamic trap chip next to a US quarter coin. C) Schematics of the active feedback mechanism. After the particle displacement was detected, the back pressure in the feedback outlet was adjusted by changing the control voltage of the electronic pressure valve. As a result, the stagnation point moved and the particle was moved by flow towards the centre of the cross slot. D) A 3D illustration of the fluidic connections showing the back-pressure control on the upper outlet. E) Vertical position (Y) trajectory of a trapped 2 µm fluorescent bead at 1.2 s^-1^ flow elongation rate in a 12.6 cP buffer. Inset: the distribution of the particle Y position. Feedback cycle time is 33 ms. F) The power density spectrum (blue) for the trace shown in (E). The black line is the nonlinear-least-square fit to a Lorentzian function, which yielded a corner frequency of 1.34 Hz. The red cross indicates the corner frequency obtained from the fitting. G) Vertical position (Y) trajectory of a trapped 2 µm fluorescent bead at 54 s^-1^ flow elongation rate. Inset: the distribution of the particle Y position. Feedback cycle time is 4.8 ms.

To measure the flow elongation rate, we flowed DNA molecules into the cross-slot with active feedback turned off. The vertical position (y) trajectory of each molecule was fit to the following equation to get the flow elongation rate *γ*_*e*_.

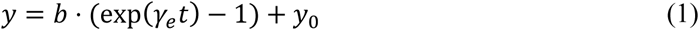

The flow elongation rate measured from ∼50 trajectories were averaged to get the final result. The flow elongation rate is a linear function of the pressure difference between the inlets and outlets. We measured the flow elongation rate at four different pressures to verify the linearity. The linear fit was then interpolated/extrapolated to the inlet/outlet pressure difference during the trapping experiment to get the flow elongation rate.

### Drag force on DNA under elongational flow

Assume dsDNA is a 2 nm diameter cylinder with length *L*. The drag force per unit length on a circular cylinder placed in a stream of uniform laminar flow along its long axis is^74^

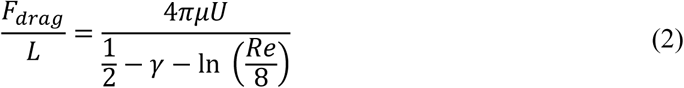

Where *U* is the flow velocity, *γ* ≈ 0.577 is the Euler’s constant, and *Re* is the Reynold’s number. Thus, the tension at *l* distance from the centre of the DNA cylinder can be calculated as

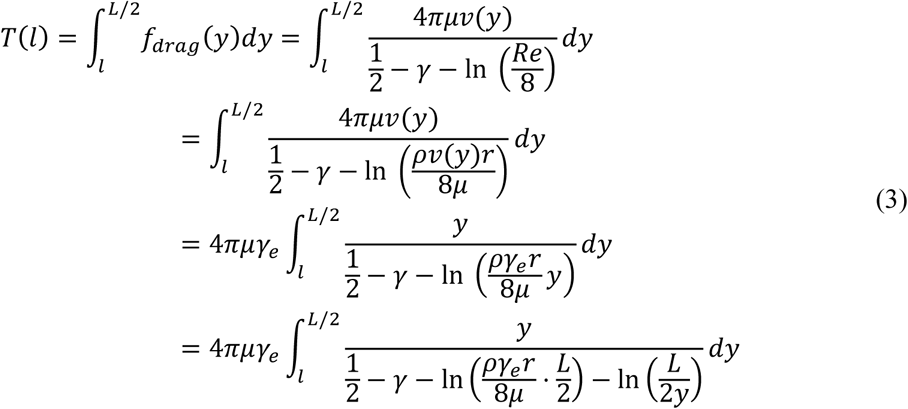

Here µ is the viscosity of the solution, *γ*_*e*_ is the flow elongation rate, *ρ* is the density of the solution, and *r* is the diameter of the rod. When 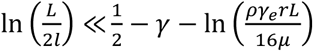, we can take the following approximation:

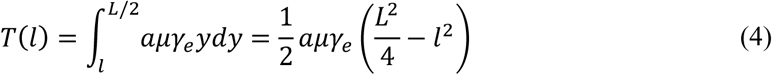

Where 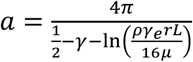.

For the 60% w/w sucrose buffer used in our experiment, *ρ* = 1.289 g ml^-1^ and *μ* = 58mPa *s*. For *γ*_*e*_ > 3 and *L* ≅ 70 μm, *a* = 0.6. 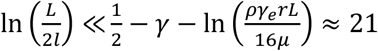. This means the above approximation holds valid for the majority of the range of *l*. For the lowest values of *l*, the fluid velocity is so low that the drag force on this segment can be neglected relative to the drag force on the rest of the filament. Therefore, Equation (4) is a good approximation for any *l*.

We note that when dsDNA stretches, the effective diameter of the filament may change—however Equation (4) suggests that this will have a negligible effect on the resulting tension. For example, let us assume that the volume of the cylinder is conserved as it is stretched. Then, the radius *r* of the cylinder would decrease by a factor of 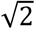 if the length were to double. According to the above calculation, such a change in *r* would have a negligible impact on the value of *a*, changing it by less than 2% for the range of parameters used in our experiments. Therefore, we can reasonably neglect the change in *r* when calculating the tension in DNA during structural transitions. This calculation also implies that small errors in our initial estimate of r will have very little quantitative effect on the calculated tension.

### Image analysis

An intensity threshold was manually selected during experiments. The threshold was selected as low as possible while ensuring no fluorescent particle would be identified when sample injection was off. When fluorescent molecules flowed into the field of view, thresholding and particle analysis were used to determine the contours of individual molecules. In order to follow the same molecule, the code looked for a molecule that was similar in position, size and brightness as compared to the molecule identified in the previous frame.

### Code availability

Custom-written Matlab scripts for data analysis are available from the corresponding author upon request, for non-profit research uses.

## Results and discussion

### High-speed active-feedback hydrodynamic trap

To apply high forces to single DNA molecules, we have developed a rigid and gas-impermeable cross-slot microfluidic chip for fluorescence-based high-speed hydrodynamic trapping. The device can trap particles under elongational flow using active feedback with direct pressure control^44^. In a cross-slot, the fluid flows in from the two inlets (the horizontal channels in Figure 1a) and exits via the two outlets orthogonal to the inlets. This creates a pure elongational field in the cross region. For a filamentous object centred in an elongational flow field, the fluid exerts drag forces against the two halves of the filament in opposite directions, inducing tension in the filament. The centre of the cross slot has zero flow velocity and is called a stagnation point. The flow elongation rate is defined as the gradient of flow velocity along the direction of flow, which in this case is the rate of velocity increase as the fluid moves away from the stagnation point. Along the inlet axis, particles are always trapped in the centre by the converging flow fields in a stable manner, but along the perpendicular outlet direction, small positional deviations from the stagnation point cause the particles to flow away with increasing velocity. Thus, the stagnation point is an unstable saddle point. However, with the addition of active-feedback, particles can be stability trapped at the stagnation point in both dimensions. As illustrated in Figure 1C, first, the position of the particle is detected by a video camera. Next, depending on the current position of the particle, the stagnation point of the flow field is moved by changing the back pressure in one of the outlets, in order to move the particle in the desired direction. The back pressure can be changed through a PDMS membrane valve in a dual-layer device^29^, or direct pressure control in a single-layer device^41^. Finally, the particle flows towards the desired trapping position (e.g. centre of the field of view) as a result of the shifted flow field. This cycle is rapidly repeated to keep the particle trapped.

To date, hydrodynamic traps have been demonstrated to trap and stretch single polymers at flow elongation rates up to 1.5 s^-1 43–46^. In order to trap and fluorescently image single molecules under high elongational flow rates, we improved feedback performance by using a rigid single-layer glass microfluidic chip (Figure 1B and D). One of the outlets is connected to a buffer reservoir filled with pressurized gas, which is controlled by an electronic pressure regulator. The other outlet is connected to a constant pressure source. Buffer injection was also driven by electronically regulated pressurized gas. This allows fast and linear control of the stagnation point position. The microfluidic chip consists of a 40 µm thick double-sided tape with channels cut-out sandwiched between a glass coverglass and a glass slide^75,76^. The tape was patterned using a die cutter^77^. Because the adhesive on the tape is deformable and the bonding between tape and glass weakens upon exposure to aqueous solution, the glass-tape-glass sandwich chip is not rigid enough for high-pressure applications. To improve the rigidity, we reinforced the chip using UV-cured optical adhesive near the channel region. We then heated up the chip to ∼100°C, which not only further improved the strength of the bonding but also smoothened the edge of channel. This rigid chip design ensures fast and stable response of the flow to pressure changes. In addition, this chip is impermeable to gas, reducing the oxygen related photobleaching and photodamage, and very durable, with no degradation in performance over a period of at least a month. These properties are desirable for not only the hydrodynamic trap but also other high-pressure and fluorescence imaging applications. Moreover, the fabrication is fast and inexpensive, requires no cleanroom equipment or special training, and therefore is accessible to almost any laboratory. Our high-speed trap allows us to trap 2 µm diameter fluorescent particles with about twice the trapping stiffness of the-state-of-art under similar conditions^41^(Figure 1E and F). We have also demonstrated stable trapping of the 2 µm fluorescent beads at flow elongation rates up to 54 s^-1^ with a moderate decrease in positional confinement (Figure 1G).

### Trapping and stretching DNA at high flow elongation rates

Using the hydrodynamic trap, we trapped and stretched YOYO-1 labelled T4 DNA to over 190% of its normal contour length in a high viscosity buffer. While DNA can be trapped and stretched in 1 cP buffer (Supplementary Figure 2), we used a high viscosity buffer, ∼ 52 cP, to maximize the drag force on the DNA. As shown in Methods, for long filaments, the drag force on each segment is proportional to the flow velocity and therefore the distance from the stagnation point (Figure 2A). As a result, the tension profile along the length of the filament is parabolic. Equation (5) shows the tension in a filament with length *L*, at a point *l* distance from the centre.

**Figure 2.**
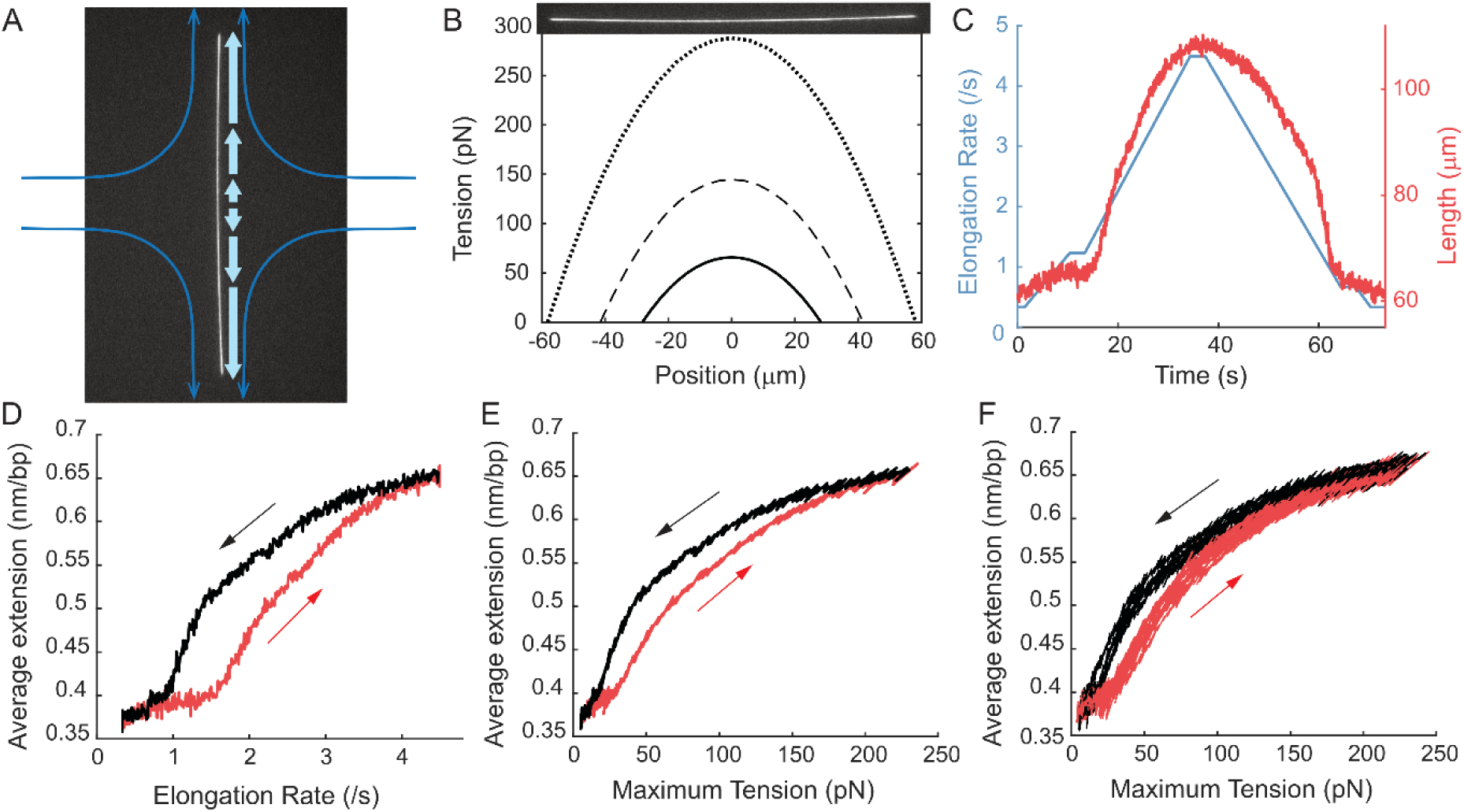
YOYO-1 stained DNA stretching under elongational flow. A) Drag force (block arrows) on each segment of a long filament in elongational flow is proportional to the distance between the segment and the stagnation point. B) Example parabolic tension profile along 56 μm (solid line), 83 μm (dashed line), and 116 μm (dotted line) long dsDNA. An image of a 116 μm polymer is overlaid on the top axis to visualize the tension distribution. C) Time traces of the flow elongation rate (blue) and DNA end-to-end distance (red). D) DNA end-to-end distance as a function of flow elongation rate. The red curve is extension and the black curve is relaxation. E) DNA end-to-end distance as a function of the maximum tension in DNA. The red curves are elongation and the black curves are relaxation. F) Overlay of results from 10 experiments. All data shown here was taken with 4:1 labelled DNA.

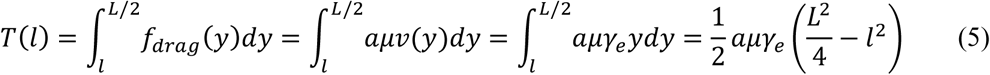

Here *a* is a constant depending on the hydrodynamic profile of the filament. Modelling dsDNA as a 2 nm diameter rod, we estimated *a* = 0.60 (Method).

The maximum tension is at the centre of the filament and proportional to the square of the filament length (Figure 2B). The tension is also proportional to the flow elongation rate and the viscosity. With a combination of a high flow elongation rate, a high viscosity buffer and long DNA, we were able to induce a maximum of 250 pN tension in DNA. We used bacteriophage T4 GT7 genome DNA with a length of 166 kbps and a contour length of 56.4 μm. We prepared DNA with two different labelling ratios, with nucleic acid base pairs to dye monomer ratios (N:D) of 4:1 and 8:1(Methods). The DNA was stained and trapped in buffer with no salt to prevent the YOYO-1 dye from dissociating. Similar results were observed in buffers with up to 20 mM NaCl. With even higher salt concentrations, the dyes dissociate too fast to allow high-speed trapping. Even though low salt conditions usually shift the equilibrium of stretched dsDNA towards melting^78^, the intercalated dyes likely protected the dsDNA from melting^15,72^. We ramped tension in the trapped DNA up and down by ramping the flow elongation rate up and down (Figure 2C, Supplementary Movie 1). The maximum tension in the DNA at each time point was calculated using the flow elongation rate and the measured length of the DNA filament (Figure 2D), and the length of DNA was plotted as a function of the maximum tension (Figure 2E). Because each intercalated dye monomer increases the contour length of dsDNA by 0.34 nm, the average extension of the intercalated B-form T4 DNA in our experiment was 0.37 nm/bp, when the maximum tension was held at ∼ 10pN. The end-to-end distance of T4 DNA increased rapidly when the maximum tension exceeded 25 pN. The slope decreased for tensions above 100 pN. At the maximum flow elongation rate we applied, the total length of the 4:1 labelled DNA was 0.65 nm/bp, over 90% longer than regular B-DNA. The extreme length indicated a DNA structure more extended than overstretched DNA and resembling the newly discovered hyperstretched DNA. Because the total end-to-end distance is the sum of many segments, each under different force, the central region of the DNA will be extended longer than the average. From the large average extension, the maximum extension in the central region likely approached the length of the sugar-phosphate backbone, which is twice the contour length of B-DNA. As described in the next section, we confirmed this by analysing the fluorescence intensity profile. During elongation-relaxation cycles, we observed significant hysteresis that could be reduced by lowering the flow ramping rates. But the hysteresis existed even at the slowest flow ramping rates we could achieve without incurring photodamage during the experiment (Figure 2E and F). Based on the current experiment, we could not distinguish whether this hysteresis was dynamic, meaning that the structural transition was slow, or static, meaning that the B and the putative hyperstretched states were bistable under our experimental conditions. Similar hysteresis was previously reported for dsDNA stretching and relaxation in the presence of intercalating molecules^79–81^. The hysteresis in those cases was attributed to the slow binding/dissociation of the intercalators. However, this cannot explain our observation because there was no free YOYO-1 dye in our experiment.

We also note that it is important to distinguish the putative hyperstretched state from DNA peeling or bubble formation, which can also dramatically increase the length of DNA by converting regions of dsDNA to single-stranded DNA (ssDNA). As discussed by Schakenraad *et. al.*^14^, combining force application with single-molecule fluorescence is key, as DNA peeling can be observed by a sudden reduction in fluorescence intensity in regions of emergent ssDNA. These were not observed under our experimental conditions, as further discussed in the next section. Additionally, within DNA force-extension curves, melting transitions and peeling transitions are typically marked by sharp increases in length during stretching and more significant hysteresis during relaxation— features that were not observed during our experiments^68,69^. It is possible that small bubbles of melted DNA occurred in the short DNA segments (about 5-14 base pairs depending on labelling ratio) between intercalated dyes. However, as detailed in the next section, two melted ssDNA strands loaded in parallel typically require much more force than what we applied to the DNA filament to be extended to the same maximum extension. Therefore, the highly extended state we observed most closely resembles the hyperstretched DNA from reference ^14^. We will refer to this putative hyperstretched state as HS^†^ DNA from now on for brevity.

### Fluorescence intensity profile indicates the force-dependent extension of DNA

In addition to measuring the length of the DNA filament as a function of flow-rate, we can also study the force-dependent properties of DNA by analysing the single-molecule fluorescence intensity profile. Because of the unique parabolic tension profile that we impose along each DNA molecule, we can sample a wide range of forces on a single DNA molecule under a single flow elongation rate. For example, the kymograph in Figure 3A shows the evolution of the intensity profile of a 4:1 labelled DNA molecule at 5 different flow elongation rates (Figure 3B, Supplementary Movie 2). At low flow elongation rates, the fluorescence intensity profile is uniform, independent of the tension (Figure 3A and D). At high flow elongation rates, the intensity remains the same in the lowest tension regions and decreases in the high-tension region (Figure 3E-H). The intensity as a function of tension are consistent across different flow elongation rates (Figure 3I). The intensity approaches a minimum for forces above 100 pN. Over the course of the experiment, the total intensity of the entire DNA molecule showed slow dissociation of the YOYO-1 dye but no clear changes correlated with changes in the length of the DNA (Figure 3C, Supplementary Figure 3). This suggests that the YOYO-1 did not dissociate, the intensity of each YOYO-1 did not change, and the dsDNA did not peel during the experiment, consistent with a direct transition into the hyperstretched state^14,15^. Therefore, the change in intensity profile is mostly due to the change in dye distribution that results from local stretching of the molecule. At low flow elongation rates, the average extension was 0.42 nm/bp (Figure 3B). As the tension increased, the length of DNA increased and so did the space between bound YOYO-1. Assuming the extension is proportional to the reciprocal of the intensity, we plotted the DNA extension as a function of the tension (Figure 3J). The results from all 5 flow elongation rates are plotted together to show the overlap between them. The black curve is the average of all the data. At low forces, around 10 pN or lower, the DNA could diffuse in and out of the focal plane, making the measured intensity lower than the actual value. Therefore, this region, to the left of the dashed line in Figure 3I, is not analysed. The extension first slightly increased from 10 to ∼25 pN, consistent with the high force region of a worm-like-chain force-extension curve^82^. This suggests that the non-intercalated segments of DNA remained in B-form in this regime. The local extension started to increase above 25 pN, consistent with the end-to-end distance measurements (Figure 2F). This increase in length indicated a gradual transition from B-DNA into HS^†^ DNA. The local extension of DNA rapidly increased to ∼0.65 nm/bp until ∼ 100 pN, and then the increase slowed down for even higher force, suggesting most of the non-intercalated base pairs of DNA had transitioned to the HS^†^ state above 100 pN. As the maximum tension increased, the region containing HS^†^-DNA expanded (Figure 3A, yellow arrows). Above 200 pN, the local extension of DNA reached ∼ 0.7 nm/bp, close to the contour length of the sugar-phosphate backbone. In comparison, two melted ssDNA strands loaded in parallel would require ∼ 500pN in total to be extended to the same length under similar no salt conditions^83^, which does not match our observation. On the other hand, our curve (Figure 3J) matches well with the DNA hyperstretching curve measured by optical tweezer experiments in the presence of the fast equilibrating dye YO-PRO-1 (Fig. S1 in reference ^15^). Both curves feature a slow increase in extension in the low-force regime, a rapid increase until ∼100 pN, and a slow increase above 100pN. The lower force regimes of both curves deviate from the usual dsDNA force-extension curves that can be described by the WLC model due to dye intercalation and hyperstretching. This suggests that our estimation of drag force on DNA is reasonably accurate. In their experiment, the gradual transition of DNA from B form to hyperstretched form was achieved through the combination of stretching and the intercalation of more dye molecules. In our experiment, as there was no free dye in solution during the stretching experiment, the non-intercalated DNA segments were transitioned to the HS^†^ form just by force.

**Figure 3.**
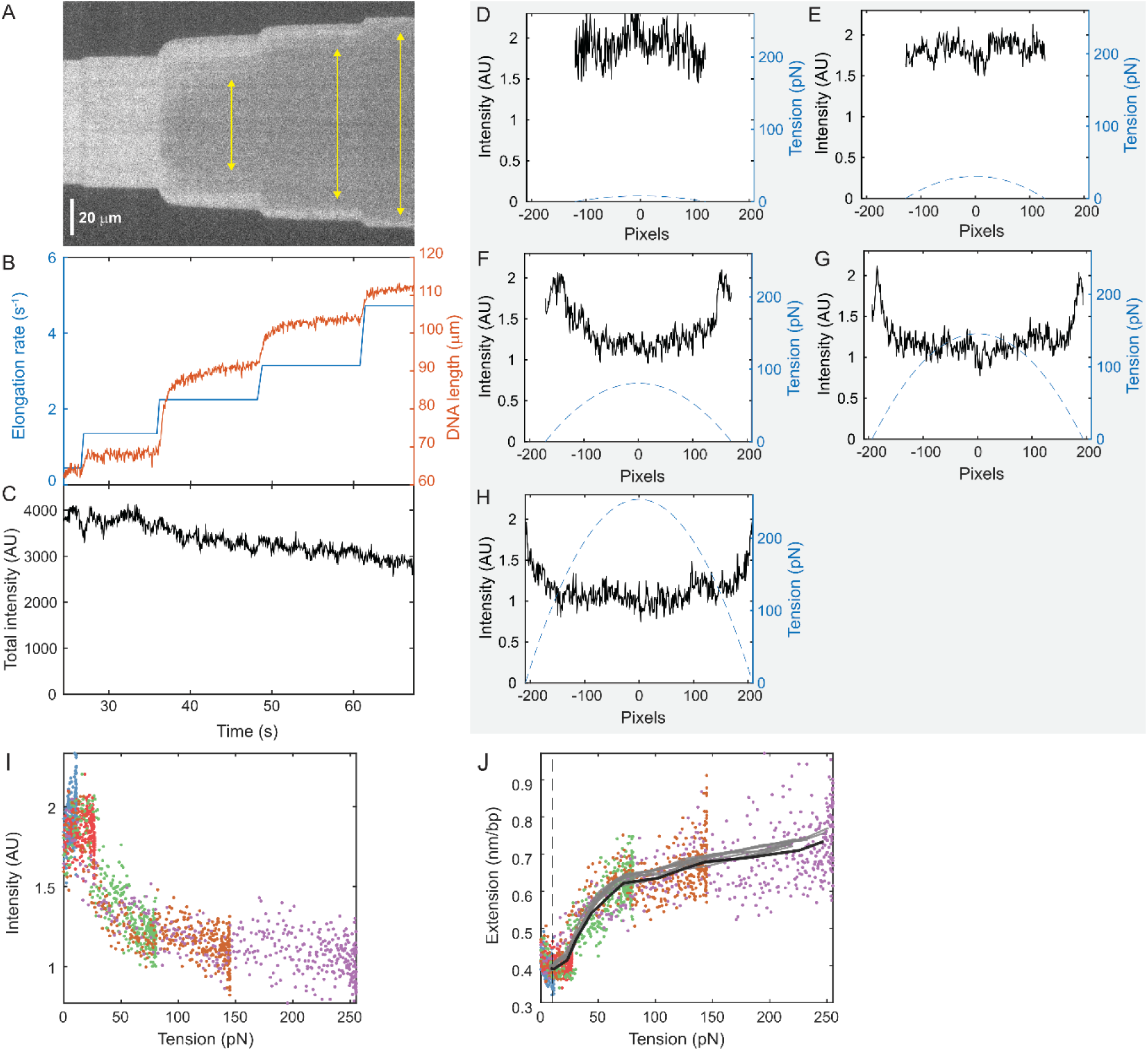
Force-dependent fluorescence intensity profile on 4:1 labelled DNA. A) Kymograph of a T4 DNA in the hydrodynamic trap as the flow elongation rate was increased in steps. Yellow arrows indicate regions of extreme extension, where the filament was extended to at least 0.6 nm/bp. B) Time traces of the flow elongation rate and the DNA end-to-end distance. Blue trace is the flow elongation rate and red trace is the end-to-end distance of DNA. C) Time trace of the total fluorescence intensity on DNA shows no correlation with the DNA length. D-H) The intensity and tension profile of the DNA at 5 different flow elongation rates. The solid curve is the intensity and the dashed line is the tension. I) The intensity as a function of tension; coloured dots are results from 5 different flow elongation rates. J) The extension per base pair of DNA as a function of tension; coloured dots are results from 5 different flow elongation rates. The black line is the average of these dots, and the gray lines are results from 10 other experiments. The region below 10 pN to the left of the dashed line was not analysed.

### Force-induced transitions to HS^†^ DNA also occur in more sparsely-labelled DNA

The transition to a state with a 0.7 nm/bp contour length was also observed in DNA with a lower staining ratio of 8:1 (Figure 4), where non-intercalated regions were longer. At this staining ratio, only approximately 12% of the DNA was intercalated. Because YOYO-1 is a dimer that intercalates into the spaces between four consecutive base pairs^84^, this corresponded to an average of 14 non-intercalated DNA base pairs between neighbouring pairs of intercalating dimers. Therefore, the majority of the DNA was not intercalated. With this lower staining ratio, the fluorescent signal was weaker, limiting the trapping to lower flow elongation rates. As a result, the maximum tension we could apply to these DNA molecules in this case was limited to approximately ∼ 150 pN. Nevertheless, the entire DNA filament could be stretched to an average distance per base pair ∼0.62 nm/bp, longer than the contour length of S-DNA. The force-dependent intensity profile of the 8:1 labelled DNA showed similar behaviour as the 4:1 labelled DNA (Figure 3). The calculated extension as a function of tension from the 8:1 labelled DNA is shown in Figure 4B, demonstrating high similarity to the results for the 4:1 labelled DNA. A direct comparison between these two conditions is presented in Figure 4C, showing an overlay of the results from the two staining ratios, with 10 replicas each. The results are very similar, though we note that the 8:1 labelled DNA molecules have a shorter extensions at low force because of their shorter contour length in the partially intercalated B-DNA form^71^.

**Figure 4.**
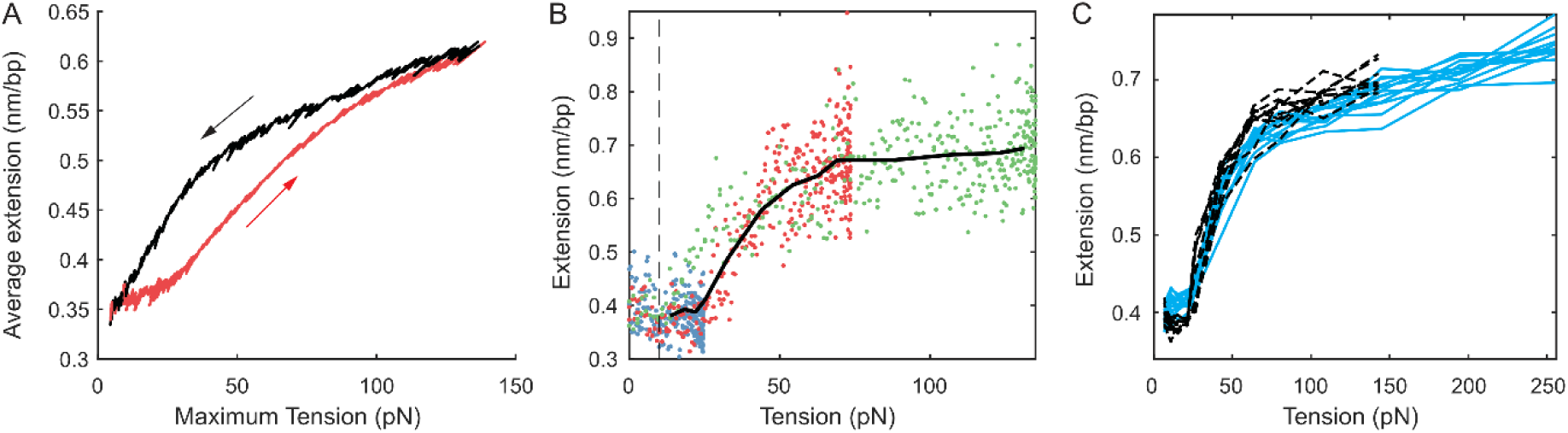
Stretching of 8:1 labelled DNA. A) Time traces of the DNA end-to-end distance as a function of the maximum tension in DNA for 8:1 labelled DNA. The red curves are elongation and the black curves are relaxation. B) The extension per base pair of DNA as a function of tension. The coloured dots are results from 3 different flow elongation rates; the black line is the average of these results. The region below 10 pN to the left of the dashed line was not analysed. C) Overlay of average force-extension curves from 20 experiments. The 10 blue lines are with 4:1 labelled DNA. The 10 black lines are with 8:1 labelled DNA.

In the previous report of hyperstretched DNA, more and more dyes intercalated in the dsDNA as the force increased. According to the model, this helped stabilizing the hyperstretched conformation through the negative binding free energy. Our observation of dsDNA reaching an extension of ∼ 0.7 nm/bp under tension with significantly fewer intercalated dyes supports the hypothesis that force alone is capable of converting dsDNA from B-form to hyperstretched form in regions up to ∼14 base pairs long. We cannot completely rule out the possibility that the intercalated dyes at the ends of these segments aided the structural transition, perhaps by disfavouring melting and overstretching transitions in the DNA segments between them^15,72^. However, it was observed that dye intercalation does not change the persistent length of B-DNA nor the orientation of the neighbouring base pairs^15,71,72,85^, which suggests intercalated dyes do not have long range impact on DNA structure.

The application of these extremely high forces to a single DNA molecule is enabled by our newly developed high-speed hydrodynamic trap with a gas-impermeable microfluidic chip. The maximum tension we induced in DNA was ∼ 250 pN, perhaps the highest constant stretching force applied to dsDNA in the literature. In principle, high forces can also be induced under lower flow elongation rates using even longer DNAs, which are not commonly commercially available. However, the ability to trap single polymers at higher flow elongation rates than before allows a wider range of DNA to be studied under a wider range of forces using hydrodynamic traps. For example, as shown in Supplementary Figure 2, the end-to-end distance of λ DNA can be measured under ramping flow elongation rates in a buffer with no added viscosity.

Our surface-free approach allows simultaneous fluorescence detection and force spectroscopy, especially in force regimes that are traditionally difficult to study. In addition, the system can be less expensive and easier to set up than many commonly used force probes, particularly for users who already have microscopes capable of fluorescence imaging. Additionally, because no bead or surface attachments are needed for this method, sample preparation can also be simplified. While traditional force spectroscopy methods, such as AFM, optical tweezer and magnetic tweezer, have advantages that include improved spatial precision and the independence of force on polymer length and buffer viscosity, the unique benefits of the hydrodynamic trapping method make it a good and complementary alternative to existing methods for certain applications.

## Conclusions

We have demonstrated force-induced stretching of dsDNA to the length of the sugar-phosphate backbone, a state that matches the newly discovered hyperstretched DNA, with a high-speed hydrodynamic trap. This experiment not only has implications for understanding the mechanics and structure of DNA under stress, but also demonstrates the potential of the hydrodynamic trap as a powerful yet accessible force spectroscopy method, able to probe molecular mechanics in previously difficult to access regimes. Not only does it enable the application of extreme forces, but it also allows biomaterials, such as cytoskeletal filaments and chromatin, to be stretched without the distortions that can be induced by surface interactions, or the need to optimize surface attachment chemistries. Furthermore, the rapid and inexpensive fabrication techniques that we have developed to produce rigid and gas impermeable microfluidic devices will be useful for a wide range of applications, especially those that require high pressure or fluorescence imaging, and will be accessible to laboratories not specialized in microfabrication.

## Acknowledgement

We acknowledge help from the microfluidic prototyping facilities in Wyss Institute for Biologically Inspired Engineering on fabricating the microfluidic chip. We are also grateful to James Pelletier, the MIT Media Lab, and the Microfluidics Core Facility at Harvard Medical for their help in fabricating early versions of the microfluidic chip. This work was supported by the National Science Foundation Graduate Research Fellowship DGE-1144152 (D.Y.), NIH NIGMS R35 GM119537 (W.P.W.), and internal support from the Wyss Institute at Harvard.

