## Supplementary Figure for "Stretching DNA to twice the normal length with single-molecule hydrodynamic trapping"

### Supplementary figures

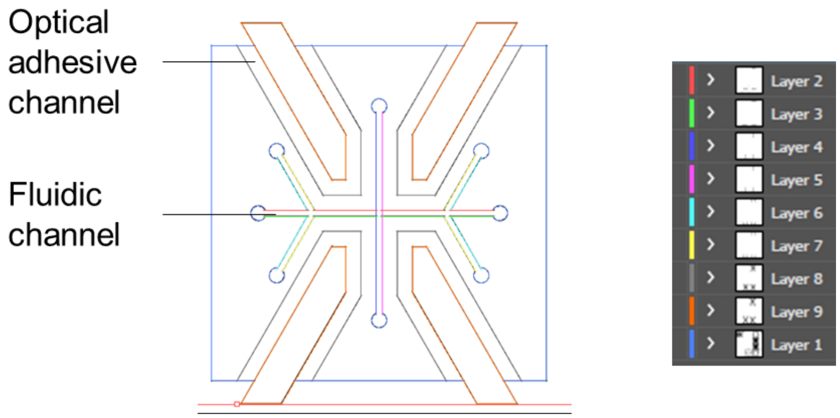


Figure 1 Design pattern of the microfluidic chip for the hydrodynamic trap. The die cutter cut the pattern following the layers listed on the right from top to bottom.

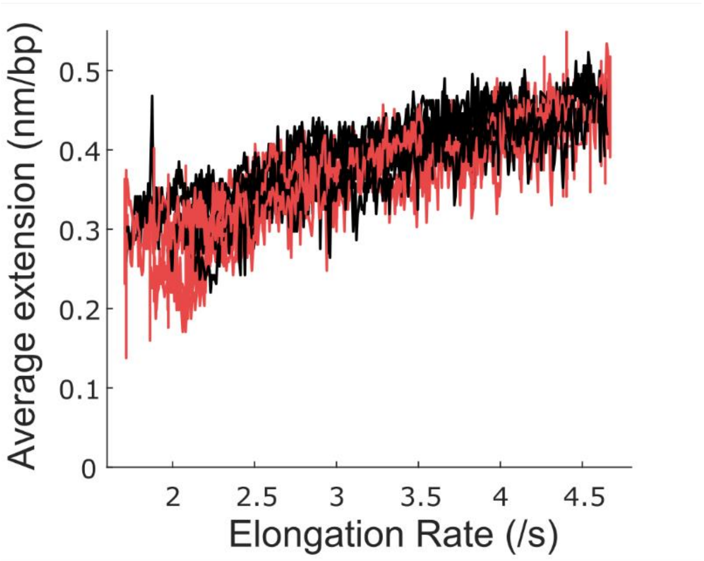

Figure 2 DNA end-to-end distance as a function of flow elongation rate. The red curve is extension and the black curve is relaxation. Data from 10 experiments are overlaid.

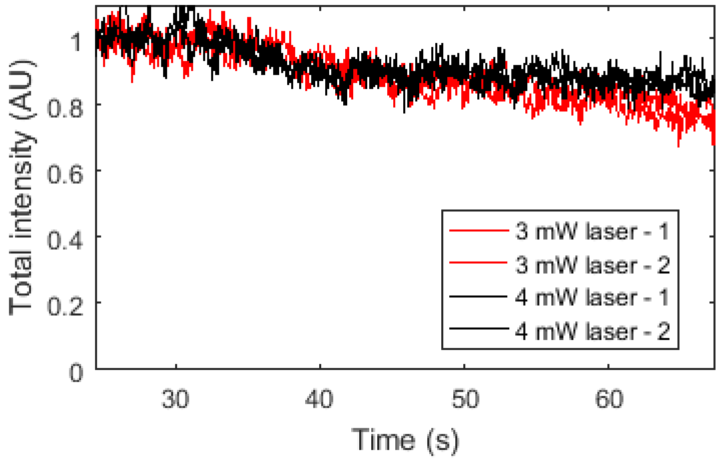


Figure 3 Time traces of the total fluorescence intensity on DNA normalized to the initial intensity at two different laser illumination powers. The traces show little dependence on illumination intensity, suggesting that the decrease in intensity is most likely due to slow dissociation of intercalating dyes.

### Supplementary movie legends

Movie 1. 4:1 labelled dsDNA stretching and relaxing under elongational flow. The data shown in Figure 2 is from this experiment. Scale bar is 20 µm.

Movie 2. 4:1 labelled dsDNA stretched using stepwise increasing elongation flow. The data shown in Figure 3 is from this experiment. Scale bar is 20 µm.
